# Pre-attentive representation of prediction certainty in autism: A mismatch negativity (MMN) study

**DOI:** 10.1101/2023.06.06.543878

**Authors:** Seydanur Reisli, Sophie Molholm

## Abstract

According to predictive processing theories of perception, the brain generates predictions to prepare for sensory input, and calibrates certainty of predictions based on their likelihood. When an input doesn’t match the prediction, an error signal leads to updating of the predictive model. Prior research suggests altered prediction certainty in autism, but predictive processing occurs across the cortical hierarchy, and the stage(s) of processing where prediction certainty breaks down is unknown. We therefore tested the integrity of prediction certainty in autism at pre-attentive and relatively automatic processing stages using the pre-attentive Mismatch Negativity (MMN) brain response. The MMN occurs in response to a “deviant” presented in a stream of “standards” and is measured while the participant performs an orthogonal task. Most critically, MMN amplitude typically varies with the level of certainty associated with the prediction. We recorded high-density EEG while presenting adolescents and young adults with and without autism with repetitive tones every half second (the standard) interspersed with infrequent pitch and inter-stimulus-interval (ISI) deviants. Pitch and ISI deviant probabilities were manipulated at 4, 8, or 16% within a block of trials to test whether MMN amplitude varied in a typical manner with respect to probability. For both groups, Pitch-MMN amplitude increased as the probability of deviance decreased. Unexpectedly, ISI-MMN amplitude did not reliably vary by probability in either group. Our Pitch-MMN findings suggest intact neural representation of pre-attentive prediction certainty in autism, addressing a critical knowledge gap in autism research. The implications of these findings are considered.

**LAY SUMMARY:** Our brains are always trying to predict what will happen next. For example, when you open your utensil drawer, it would be surprising to see books because your brain expected to see utensils. In our study, we looked at whether the brains of autistic individuals automatically and accurately recognize when something unexpected happens. Results showed similar brain patterns in individuals with and without autism, suggesting that responses to prediction violations are generated in a typical manner during early cortical information processing.

## INTRODUCTION

A key brain function guiding adaptive behavior is the ability to calibrate the confidence of predictions in the face of uncertainty (Bar et al., 2006; Feldman & Friston, 2010). For example, if your flight is departing from an airport that you use for the first time, estimating the time it would take to make your way to the airport and get through security would be difficult, so you would give yourself ample “extra” time, and you may plan to arrive at the airport. However, in cases where information guiding a prediction is more reliable, such as when you are flying from an airport that you often use and know well, predictions can be made more confidently. Therefore, predictions are probabilistic by nature and can be associated with more or less certainty, dependent on the reliability of the given information (Clark, 2015; Friston, 2012). It is essential to estimate the probability that a prediction is correct, and to assign a proper level of confidence in order to deal with uncertainty in the environment effectively.

Recent studies suggest that the representation of prediction certainty (i.e., precision) is altered in autism (Lawson et al., 2014; Palmer et al., 2017). Furthermore, classic autistic symptoms have been attributed to altered predictive processing. For example, a reduced ability to tune the certainty of predictions could prevent distinguishing informative cues from non-informative ones in the complex and rich nature of social life for autistic individuals, leading to social difficulties (Pellicano & Burr, 2012; Van de Cruys et al., 2014), and resistance to change was linked to discomfort upon changes due to impaired predictive processing (Gomot & Wicker, 2012).

A number of studies suggest that autistic individuals have problems adjusting the certainty of predictions based on changing environmental statistics (Chambon et al., 2017; Coll et al., 2020; Spee et al., 2022), or using context in adjusting the certainty of predictions (Arthur et al., 2020; Randeniya et al., 2021; Sapey-Triomphe et al., 2021). Importantly, in these studies, the integrity of predictive processing in autism was assessed using active tasks. Given that the human brain generates predictions automatically even without the individual’s awareness (Bar, 2007; Clark, 2013), it remains unclear whether the observed problems stem from issues in primitive stages of predictive processing such as automatic detection and registration of stimulus probabilities, or systems that rely on attention and higher cognitive processes.

To investigate the pre-attentive calibration of prediction certainty in the autistic brain, we leveraged a well-established brain potential called the mismatch negativity (MMN) that can be measured from the human scalp using electroencephalography (EEG) (Näätänen, 1995; Näätänen et al., 2007). The MMN is generated when a repeated stimulus (a standard) is replaced with a different stimulus (a deviant). It is commonly considered to index that the brain has registered a deviation from the sensory memory trace of the repeated stimulus (Molholm et al., 2005; Winkler, 2007). In the framework of predictive coding, the MMN represents a pre-attentive prediction error response (Fitzgerald & Todd, 2020; Knight et al., 2020). The amplitude of the MMN increases as the deviant probability decreases (Evstigneeva & Aleksandrov, 2009; Fisher et al., 2011; Haenschel et al., 2005; Sabri & Campbell, 2001). It is theorized that this is because the strength of a memory trace is greater for more frequently presented standards, and accordingly, larger MMN responses to deviants are elicited due to the violation of more robust (e.g., certain) predictions (SanMiguel et al., 2021; Sato et al., 2000; Tsogli et al., 2019; Winkler & Czigler, 1998). Therefore, the MMN provides a powerful tool to assess the pre-attentive representation of the level of certainty associated with predictions.

To test whether MMN amplitude varies with respect to deviant probability in autism, we presented three levels of deviant probability (4%, 8%, and 16%) while measuring EEG activity in young adults with and without autism. We presented both pitch and inter-stimulus-interval (ISI) deviants. Pitch deviants are commonly used in auditory MMN studies, whereas the addition of an ISI deviant allowed us to control for potential neural refractory effects caused by changes in the physical properties of the stimulus (Jacobsen & Schröger, 2001). Altogether, the study fills a critical knowledge gap in the autism literature by addressing whether prediction certainty is flexibly calibrated in pre-attentive stages of cortical information processing in the autistic brain.

## METHODS

### Participants

Eighteen autistic individuals (age: 21.3 ± 3.6 years old; 14 males; Intelligence Quotient (IQ): 104.9 ± 19.3) and 15 IQ- and age-matched control subjects (age: 20.7 ±3.2 years old; 8 males; IQ: 104.1 ± 12.5) participated in the study, all with normal hearing, aged between 17 and 29 years. Participants were recruited without regard to sex, race, or ethnicity.

For both groups, exclusion criteria included a performance IQ below 80; a history of head trauma; premature birth; a current psychiatric diagnosis; or a known genetic syndrome. For the control group, having a biological first-degree relative with an autism diagnosis or having a history of developmental, psychiatric, or learning disorders were additional exclusion criteria. Participants who were on stimulant medications for attention-deficit/hyperactivity disorder were asked to not take them at least 24 hours prior to the experiment. All components of the study were approved by the Institutional Review Board at Albert Einstein College of Medicine. Written informed consent was obtained prior to conducting the research from participants for participants ages 18 or older. For the participants who were younger than 18, written consent was collected from their legal guardians in addition to obtaining written assent from themselves.

### Neuropsychological and clinical testing

IQ was estimated using the Wechsler Abbreviated Scale of Intelligence (Simard et al., 2015). For the autism group, an autism diagnosis was confirmed with the Autism Diagnostic Observation Schedule, Second Edition (ADOS-2) (Lord et al., 2012), the Autism Diagnostic Interview-R (Lord et al., 1994), and expert clinical judgment by a licensed psychologist at the Human Clinical Phenotyping Core of Albert Einstein College of Medicine Rose F. Kennedy Intellectual and Developmental Disability Research Center.

### Stimuli and task

Participants were presented with frequently and infrequently occurring auditory tones (standards and deviants, respectively) in a passive oddball design while they watched a movie of their choice with the sound off and subtitles on. Standards were 100 ms tones of 1000 or 1300 Hz presented at an inter-stimulus interval (ISI) of 500 ms. Two types of deviants, frequency (1000 or 1300 Hz, depending on the frequency of the standard) and ISI (300 ms) (Fig. 1) were presented within the same block, and the probability of a given deviant was varied across blocks between 4%, 8%, and 16%. Because the MMN system is considered to generate predictions at the individual feature level (Molholm et al., 2005), the probability of the standard representation of ISI or Pitch was considered to be the inverse of the ISI or Pitch deviance. Piloting confirmed that robust and similar MMNs were elicited when the two deviants were presented in the same block versus in different blocks. The order of deviants and standards was pseudo-randomized within each block so that a standard always preceded and followed a deviant.

**FIGURE 1:**
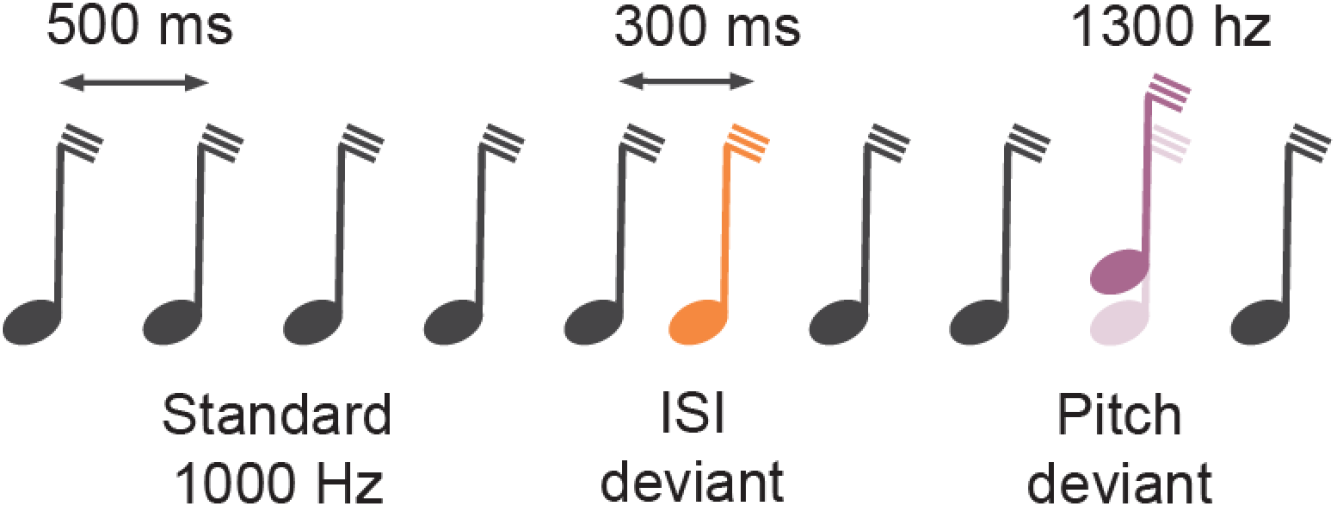
Schematic of experimental paradigm and probability conditions. Standard (black) tones are delivered at every 500 ms. ISI deviants are presented 300 ms following the standard. Pitch deviants were 1300 Hz tones among 1000 Hz standards or vice-versa.

The experiment consisted of two sessions, with a ∼15-min break in between. For the majority of subjects, a total of 8 blocks, each consisting of ∼1200 stimuli, were presented in the order of 4-8-16-4% in the first session and 4-16-8-4% in the second. Since a 4% block has 2-4 times fewer deviants than 8-16% blocks, we presented the 4% condition blocks more often, ensuring a minimum of 150 trials per participant for that condition. For some participants, the passive oddball paradigm was followed by an active oddball task, not reported here.

To ensure that analysis of the MMN is based on responses to physically identical stimuli, the frequencies of the standard and deviant stimuli were interchanged across sessions within a participant such that in session 1, the 1000 Hz tone served as the standard and 1300 Hz tone served as the deviant, while the opposite was true for session 2.

### Stimulus delivery and EEG recording

Participants were seated in a dimly lit, electrically shielded, and sound-attenuated room. They were presented with auditory stimuli delivered via in-ear headphones with disposable foam ear plugs while viewing a silent movie of their choice with subtitles on. Continuous EEG was recorded from 160 scalp electrodes at a rate of 512 Hz using the BioSemi ActiveTwo system (Amsterdam, Netherlands) referenced to a common mode sense (CMS) active electrode and a driven right leg (DRL) passive electrode. The BioSemi ActiveTwo system replaces ground electrodes that are used in conventional EEG system with CMS and DRL by creating a feedback loop, thus rendering them references.

### EEG pre-processing

EEG data were pre-processed using eeglab (Delorme & Makeig, 2004) library in Matlab and custom scripts. To reduce computing demands in subsequent analyses, we downsampled data to 128 Hz and selected a subset of 64 channels corresponding to the BioSemi 64-channel cap. EEG was digitally filtered with a high-pass filter at 1 Hz to achieve an optimal signal-to-noise ratio in subsequent ICA analyses (Winkler et al., 2015), and a low-pass filter at 45 Hz to remove line noise. A visual inspection was made per subject to look for sections of the continuous data with excessive noise (such as those that are initiated when a participant moves excessively) and bad scalp channels. On average, this resulted in the removal of two channels and 5% of time points per participant. Infomax Independent Component Analysis (ICA) was used to remove potential non-brain related activity, mainly eye-movement related muscle artifacts. For each Independent Component (IC), the iclabel program (Pion-Tonachini et al., 2019) was used to calculate the probabilities for that IC belonging to the seven different IC categories, including Brain, Muscle Noise, Eye Noise, Heart Noise, Line Noise, Channel Noise, and Other. A noise metric was created via summation of muscle-, eye-, and channel-related noise probabilities. An IC was excluded only if it had more than a 25% chance for the noise category while having less than a 25% chance of the brain category. The channels that were rejected prior to ICA were interpolated using the spherical interpolation method (Perrin et al., 1989) via the pop_interp function of eeglab. Data epochs time-locked to stimulus presentation were created for an interval starting from 50 ms pre-stimulus to 450 ms post-stimulus onset. The -50 to 0 ms time period was used for baseline correction as used in earlier MMN studies (Chen et al., 2014; Hamilton et al., 2018; Murphy et al., 2013; Rudolph et al., 2015; Rüsseler et al., 2001; Todd et al., 2014).

A basic trial rejection protocol was applied, rejecting trials below or above a threshold. This threshold was calculated separately for each subject by taking the two standard deviations from the mean of maximum points of all channels & time points for all trials, after the removal of time points exceeding 150 uV. These steps yielded rejection of 4.4% of all trials on average per subject. Following trial rejection, data were rereferenced to PO8, a channel approximating the right mastoid. This referencing scheme maximizes the MMN over the fronto-central scalp. For each trial type, an average was taken first for each subject for use in statistical analysis and across-subjects averaging within each group for data representation.

### Statistical analyses and visualization

To measure probability effects on the MMN, deviant minus standard difference waveforms were generated for each participant for each probability condition. The average amplitude from four frontocentral scalp channels (Fz, FCz, FC1, FC2); in line with the typically frontocentral foci of the MMN (Fisher et al., 2011; Goris et al., 2022; Ritter et al., 2006; Sato et al., 2000) from the 100 to 200 ms window, where the MMN was expected to be greatest (Akatsuka et al., 2005; Kekoni et al., 1997; Phillips et al., 2015; Sams et al., 1985; Todorovic & de Lange, 2012), was used for subsequent statistical analyses. For each MMN type (pitch and ISI), a separate two-way analysis of variance (ANOVA) was performed with factors of deviant probability (3 levels: 4% 8% 16%) and group (2 levels: control and autism) to test the effects of these factors on MMN amplitude. To test for the presence of significant linear relationships between deviant probability and MMN response, linear least-squares regression analyses were performed between probability condition and MMN amplitude for each deviant type and group.

## RESULTS

We designed an oddball paradigm where the probability of deviant stimuli was parametrically manipulated at three levels for each of Pitch and ISI: 4%, 8%, and 16%. We assessed the effects of deviant probability and group (control and autism) on the MMN brain response to investigate whether stimulus probability is flexibly represented in the autistic brain.

Examination of the MMN waveforms suggested that robust MMNs were elicited by all probability conditions for both groups of participants. Both autism and control groups showed typical-looking MMN responses focused over the frontocentral scalp region (Fig. 2A, Fig. 3A, also see Fig. S1 for representative standard and deviant waveforms). The MMN had frontal negativity (Fig. 2C, Fig. 3C) that peaked at ∼150 ms from stimulus onset. For the Pitch-MMN, amplitude increased as deviant probability decreased in both groups of participants (Fig. 2). The ANOVA yielded significant main effects of probability [F(2,96) = 5.632, p=0.005] and group [F(1,96) = 7.756, p=0.006], while showing no significant and group-by-condition interaction [F(2,96) = 0.033, p=0.967]. Linear least-squares regression between probability and MMN amplitude showed that Pitch-MMN amplitude was significantly more negative as deviant probability decreased for both the control (ß(45)=0.78 ± 0.33, p=0.02) and the autism groups (ß(57)=0.89 ± 0.42, p=0.04) (Fig. 2).

**FIGURE 2:**
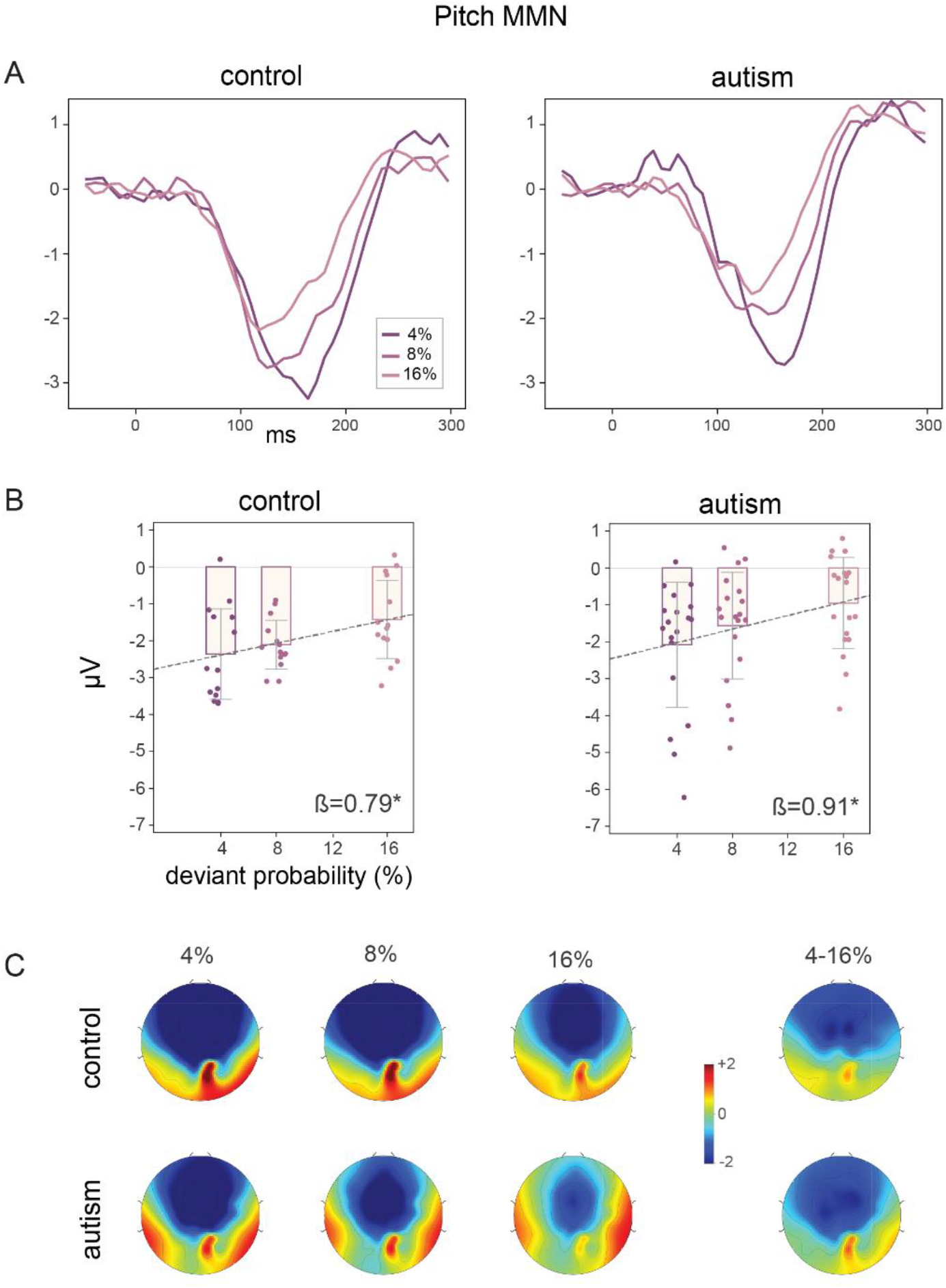
Pitch-MMNs. **(A)** MMN waveforms over the frontocentral scalp (average of Fz, FCz, FC1, FC2), calculated by subtracting the responses to standards from the response to deviants, shown for all probability conditions. **(B)** Individual subject Pitch-MMN amplitudes (calculated for the 100-200 ms window) for the 4%, 8%, and 16% probability conditions. The dotted lines show linear regression between probability and the Pitch-MMN amplitude. Slopes of the linear regression lines are shown at the bottom of the plots. Error bars show 95% confidence intervals. * denotes p <0.05. **(C)** Topographies for the 100-200 ms time interval for Pitch-MMNs at 4% and 16% deviant probability conditions and the difference topographies between 4% and 16% conditions.

**FIGURE 3:**
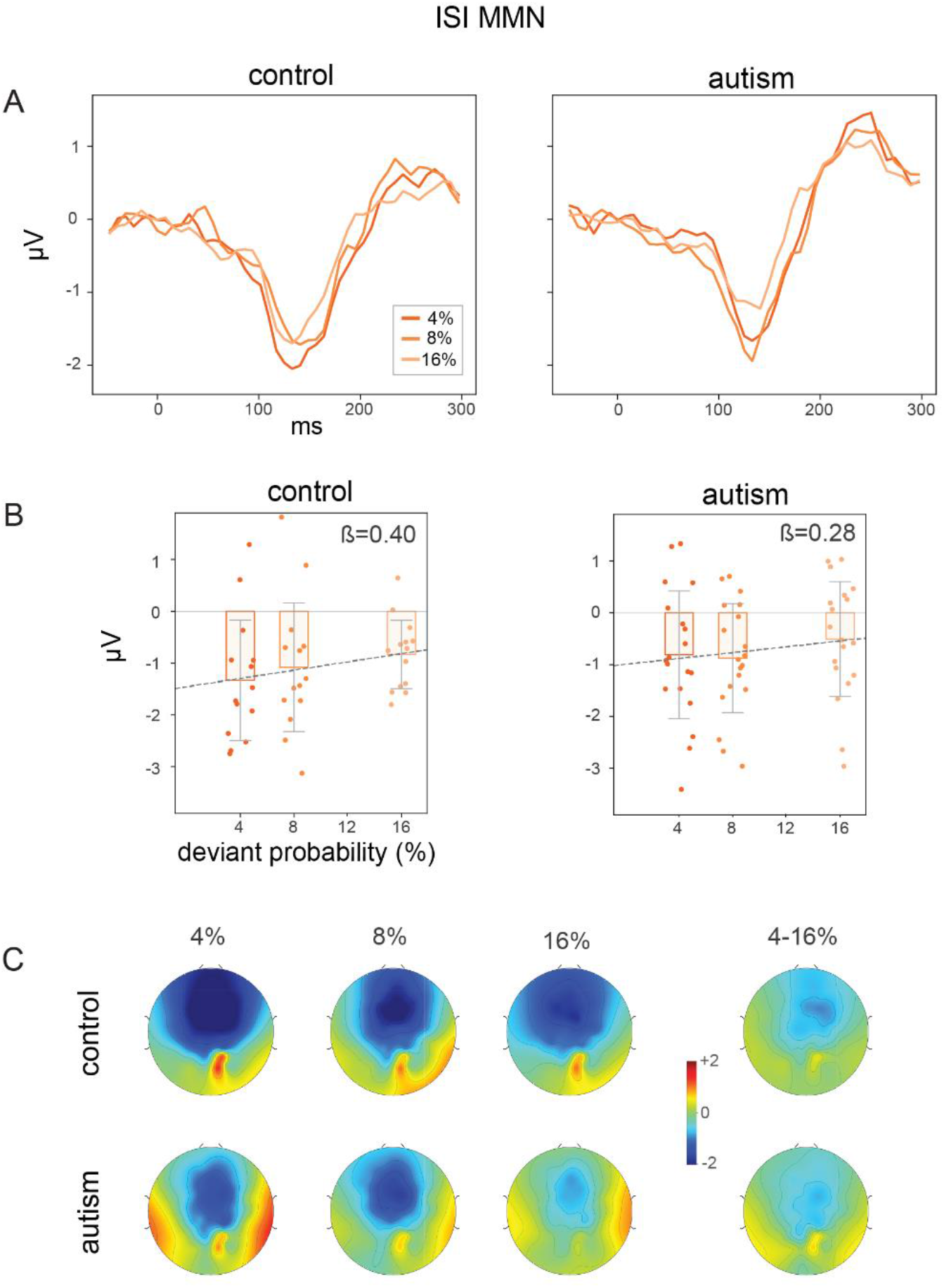
ISI-MMNs. **(A)** MMN waveforms for the ISI deviant over frontocentral scalp (average of Fz, FCz, FC1, FC2), calculated by taking the difference between the response to standards and ISI deviants separately for each probability condition. **(B)** Individual subject ISI-MMN amplitudes (calculated for the 100-200 ms window) for the 4%, 8%, and 16% probability conditions. The dotted lines show linear regression between probability and the ISI-MMN amplitude. Slopes of the linear regression lines are shown on top of the plots. Error bars show 95% confidence intervals. **(C)** Topographies for the 100-200 ms time interval for ISI-MMNs at 4% and 16% deviant probability conditions and the difference topographies between 4% and 16% conditions.

For the ISI-MMN, although the lowest probability condition yielded a numerically larger MMN than the highest probability condition in both groups (Fig. 3), the influence of deviance probability was not as visually evident. This was supported by statistical analysis showing a significant effect of group [F(1,96) = 3.998, p=0.048] but no significant effect of probability [F(2,96) = 1.168, p=0.316] or group-by-probability interaction [F(2,96) = 0.22, p=0.803]. Linear regression between deviant probability and MMN amplitude showed no significant linear relationsjip between the deviant probability and MMN amplitude for the control group (ß(45)=0.41 ± 0.34, p=0.23) and for the autism group (ß(57)=0.26 ± 0.32, p=0.41) (Fig. 3).

**SUPPLEMENTARY FIGURE 1:**
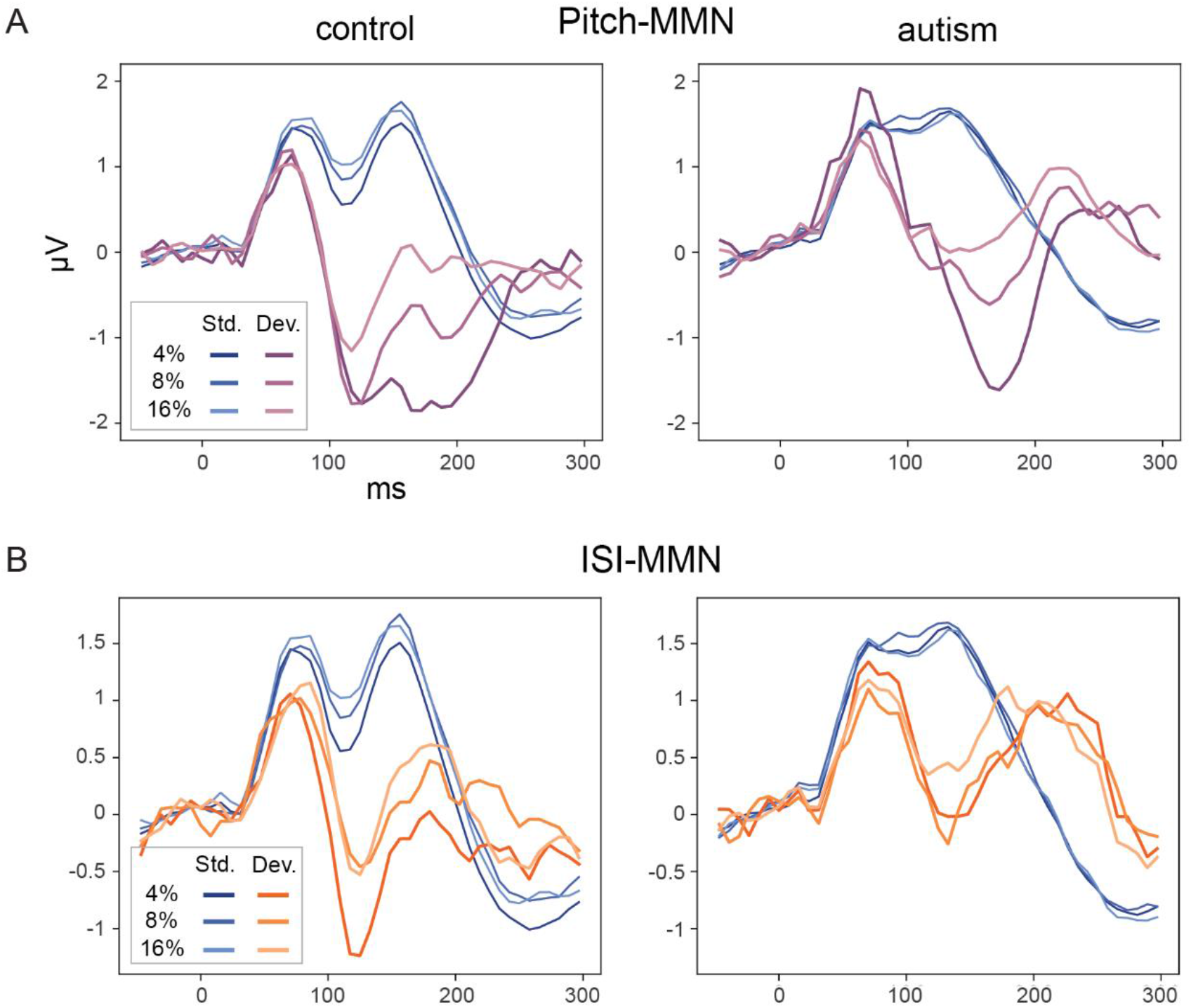
Standard and Deviant Waveforms. **(A)** ERP waveforms to pitch deviants (purple shades) and standards (blue shades) over the frontocentral scalp (average of Fz, FCz, FC1, FC2). **(B)** ERP waveforms to ISI deviants (orange shades) and standards (blue shades) over the same scalp region as A.

## DISCUSSION

Here we show evidence for intact pre-attentive processing of prediction certainty in autistic individuals. In particular, the influence of probability on the MMN was present in autistic individuals and did not differ from the probability effects in the control group.

While such probability effects are a robust finding in the MMN literature (Akatsuka et al., 2007; Deacon et al., 1998; Javitt et al., 1998; López-Caballero et al., 2016; Molholm et al., 2004), to our knowledge this is the first time that probability effects on the MMN have been examined in autism. The finding of intact probability effects on the MMN in autism is important as it suggests that fundamental neural mechanisms underlying pre-attentive evaluation of certainty of predictions are preserved in this population, potentially paving the way for further research into the post-attentive stages of predictive processing in autism and the search for targeted interventions accordingly.

The present study contributes to the literature on predictive processing in autism by specifically addressing the question of whether atypical weighting of prediction errors suggested by previous studies might reflect impairment at early pre-attentive stages of processing. Several prediction coding based theories of autism have suggested that autistic individuals do not represent prediction errors (deviation from expectations) in a flexible manner (Friston, 2013; Lawson et al., 2014; Palmer et al., 2015; Van Boxtel & Lu, 2013; Van de Cruys et al., 2014). However, studies supporting this idea have used active prediction tasks (Arthur et al., 2021; Lawson et al., 2014; Reisli et al., 2023), and it remained unclear whether these difficulties arose from deficits in pre-attentive stages of processing of prediction certainty, or from an inability to actively use this information in performing a task. The current study provides evidence to suggest that the latter may be the case. Thus the autistic brain is able to represent the likelihood of events pre-attentively but may struggle with using this statistical information to modulate certainty of predictions at a later stage of information processing (Garrido et al., 2009; Wacongne et al., 2012).

An unexpected finding from the present study is that probability effects were not present for the ISI-MMNs, for either group. Although examination of the mean waveforms from each of the groups suggested the presence of probability effects in the predicted direction (the MMN was numerically larger for the condition in which deviant probability was lowest compared to when it was highest), this was not statistically significant. The literature on probability effects on the MMN has largely involved pitch deviants and does not provide a basis on which to speculate on reasons for the lack of ISI-MMN modulation by probability in the current study. However, given that a previous study using larger probability differences showed probability effects on the ISI-MMN (Deacon et al., 1998), one possibility is that probability effects on ISI-MMNs are only reliably present for larger probability contrasts. This, unfortunately, only raises the question of just why the sensitivity of representation of the likelihood of different stimulus features (e.g., pitch versus ISI) would vary, a question that, while of interest, is beyond the scope of the current investigation.

Finally, it is important to note that the MMN amplitude was significantly smaller in the autism group compared to the control group (for both Pitch-MMN and ISI-MMN). This is in line with previous reports suggesting reduced MMN in autism (Vlaskamp et al., 2017; Lassen et al., 2022, Abdeltawwab et al. 2015), although other MMN studies report no differences or even increased MMN amplitude (Bonnel et al., 2003; Knight et al., 2020). The reason for this across-study variability is not well understood, although it may be linked to the specific features being studied or represent individual variability in the samples across different studies.

In conclusion, our results suggest that the representation of prediction certainty is intact in autistic individuals at pre-attentive stages of information processing. Achieving a comprehensive understanding of predictive processing in autism is important for informing the development of targeted therapies, such as the creation of cognitive-behavioral theories for responding to unpredicted changes in the environment.

## ACKNOWLEDGEMENTS

We are grateful to the individuals who participated in this research and their families for their time and their commitment to the advancement of scientific discovery; without them, this work would not be possible. We thank Dr. Catherine Sancimino and Dr. Juliana Bates, who administered or supervised the clinical assessments, and Sarah Barkley for her help in setting up the paradigm as a summer intern in the lab. We are also grateful to the research assistants and technicians at the Cognitive Neurophysiology lab of Albert Einstein College of Medicine who contributed to the collection of high-quality EEG data. The Human Clinical Phenotyping Core, where the participants enrolled in this study were clinically evaluated, is a facility of the Rose F. Kennedy Intellectual and Developmental Disabilities Research Center (IDDRC), which is funded through a center grant from the Eunice Kennedy Shriver National Institute of Child Health & Human Development (NICHD U54 HD090260; P50 HD105352).

